# Proprioception Has Limited Influence on Tactile Reference Frame Selection

**DOI:** 10.1101/2020.11.02.364752

**Authors:** Or Yizhar, Galit Buchs, Benedetta Heimler, Doron Friedman, Amir Amedi

**Affiliations:** Department of Cognitive Sciences, The Hebrew University of Jerusalem, Israel; Baruch Ivcher School of Psychology, Interdisciplinary Center Herzliya, Israel; Sammy Ofer School of Communications, Interdisciplinary Center Herzliya, Israel

## Abstract

Perceiving the spatial location and physical dimensions of objects that we touch is crucial for goal-directed actions. To achieve this, our brain transforms skin-based coordinates into a reference frame by integrating visual and proprioceptive cues, a process known as tactile remapping. In the current study, we examine the role of proprioception in the remapping process when information from the more dominant visual modality is withheld. We developed a new visual-to-touch sensory substitution device and asked participants to perform a spatial localization task in three different arm postures that included posture switches between blocks of trials. We observed that in the absence of visual information novel proprioceptive inputs can be overridden after switching postures. This behavior demonstrates effective top-down modulations of proprioception and points to the unequal contribution of different sensory modalities to tactile remapping.

## INTRODUCTION

How does body posture influence the way we interpret and perceive the external environment? Our constant physical interaction with the world requires a continuous update of the body’s location in external space, its relation to other objects, and its relation to itself (e.g., the relative positions of body-parts in motion). These varied representations of the body form a conscious perception of the external world and play an essential role in action planning. Think for instance of driving a car with a stirring wheel in your hand - sensory information about the wheel gives rise to a coherent perception of its function and leads to a set of possible actions that one can perform with it. First, we access the wheel’s physical dimensions through tactile stimulations received on our palms that form an anatomical reference frame. To steer the car, we transfer this information into a different reference frame that integrates the anatomical reference frame with external information fitting the wheel’s functional use (e.g. the sidewalk is to *right*, the opposite lane is to *left*), in a process known as tactile remapping^1,2^. The transformation results in the adoption of an allocentric reference frame that is independent of the body, relating objects’ dimensions to external anchors, or of an egocentric reference frame, relating objects’ positions to one’s own body^3,4^. Tactile stimulations can be remapped into many allocentric or egocentric reference frames with the ultimate selection depending on the actions that precede or follows the sensation^5,6^, the gravitational dimensions of the external environment ^7–10^, and the general position of the body^2,4,11^.

Which cognitive mechanisms drive tactile remapping? One influential view considers tactile remapping as part of a wider process of acquiring sensorimotor contingencies^2,8,12,13^. According to this theory, perception emerges through experiencing multiple co-patterns of incoming sensory signals coupled with outgoing motor actions towards the stimulus. In the context of tactile remapping, multiple reference frames are learned from exposure to tactile stimulations that are integrated with visual and proprioceptive cues to execute diverse actions. Thereafter, many reference frames are accessible with different probability weights that are constantly updated with ongoing sensorimotor experiences, and are then retrieved implicitly during the tactile remapping process^2,3,13^. Supporting studies show that a change to body posture, gaze, or object’s position in external space triggers a gradual adaptation period marked by inconsistent reference frame selections and increases in decision times as participants integrate new sensory information^4,10,14,15^. Over time, participants’ reference frames selection becomes more robust as new contingencies are established^2,10,16,17^.

However, the description of tactile remapping as a byproduct of sensorimotor contingencies overlooks the differential contribution of vision and proprioception to the process. In particular, it is known that vision is our predominant sensory modality for spatial processing and has a particularly strong influence on reference frame selections. Vision impairs the localization of tactile stimuli when arms are crossed^17,18^, determines the assignment of vertical and horizontal axes to our body^14,19^, and biases actions towards touched objects^20,21^. Conversely, the effects of proprioception (i.e., posture) per se on tactile remapping are less studied and harder to isolate, even when visual inputs are withheld. Such studies typically include complex spatial and cognitive tasks that strongly influence perception but are unrelated to proprioception proper, such as the need to relocate the object after changing postures^2,5,13^, or a manual delivery of tactile stimuli that biases participants’ responses^4,14,15^.

In the current study, we tested the effects of changing body postures on reference frame selection in blindfolded participants. To disentangle the contribution of proprioception from other factors, we built a visual-to-tactile Sensory Substitution Device (SSD)^22,23^ that transforms 2D grayscale images into tactile stimuli delivered on the inner arm. The fixed device moves together with the arm and thus nullifies the need to actively relocate the stimulations. With this unique setup, we had participants perform a simple spatial localization/orientation task while also switching their arm’s posture between trial blocks. We aimed to further investigate the role of posture in tactile remapping by asking how switching postures affect previously acquired references. According to a strictly sensorimotor prediction, after switching postures new proprioceptive cues will gradually integrate with a stored body representation that will produce an adaptation and learning period, characterized by less consistent responses and longer decision times. Results in this direction will suggest that, albeit vision’s dominant influence, incoming proprioceptive signals strongly influence the remapping process. An alternative hypothesis is that proprioception is a unique sensory modality, which we are less consciously aware of ^24^, and its effect on remapping is weaker than previously assumed. In this case, participants would rapidly adapt to new postures as top-down representations would override incoming bottom-up proprioceptive cues, diverging from a pure sensorimotor contingency description.

## RESULTS

In this study, we investigated the properties of reference frames’ selection when relying solely on proprioceptive cues. To this aim, we used a visual-to-tactile SSD that transfers 2D images to blindfolded participants’ arm, which was placed in three different postures (Fig. 1a.). In each experimental trial they were presented with a series of vibrotactile stimulations corresponding to pixels an image and their task was to report the stimulus’ spatial location (“up/down”) or orientation (“upward/downward”). We interpreted their responses on the y-axis based on the anatomy of the inner arm, referred to here as coordinate selection. Distal responses were defined as a perception of the line’s upper location/orientation located away from the trunk and towards the wrist. Proximal responses were defined as the perception of the line’s upper location/orientation located towards the trunk and the elbow (Fig. 1b.). This was done to obtain a homogenous categorization of participants’ responses, ultimately facilitating the identification of their selected reference frame (arm-based, trunk-based, external, etc.). Group-level analyses were performed with a Wilcoxson signed-rank two-tailed test and corrected for multiple comparisons (Bonferroni, p = 0.05). Subject-level statistics passed the criteria for normal approximation and analyzed with a normalized two-tailed t-test corrected for multiple comparisons (False Discovery Rate, α = 0.05).

**Fig 1.**
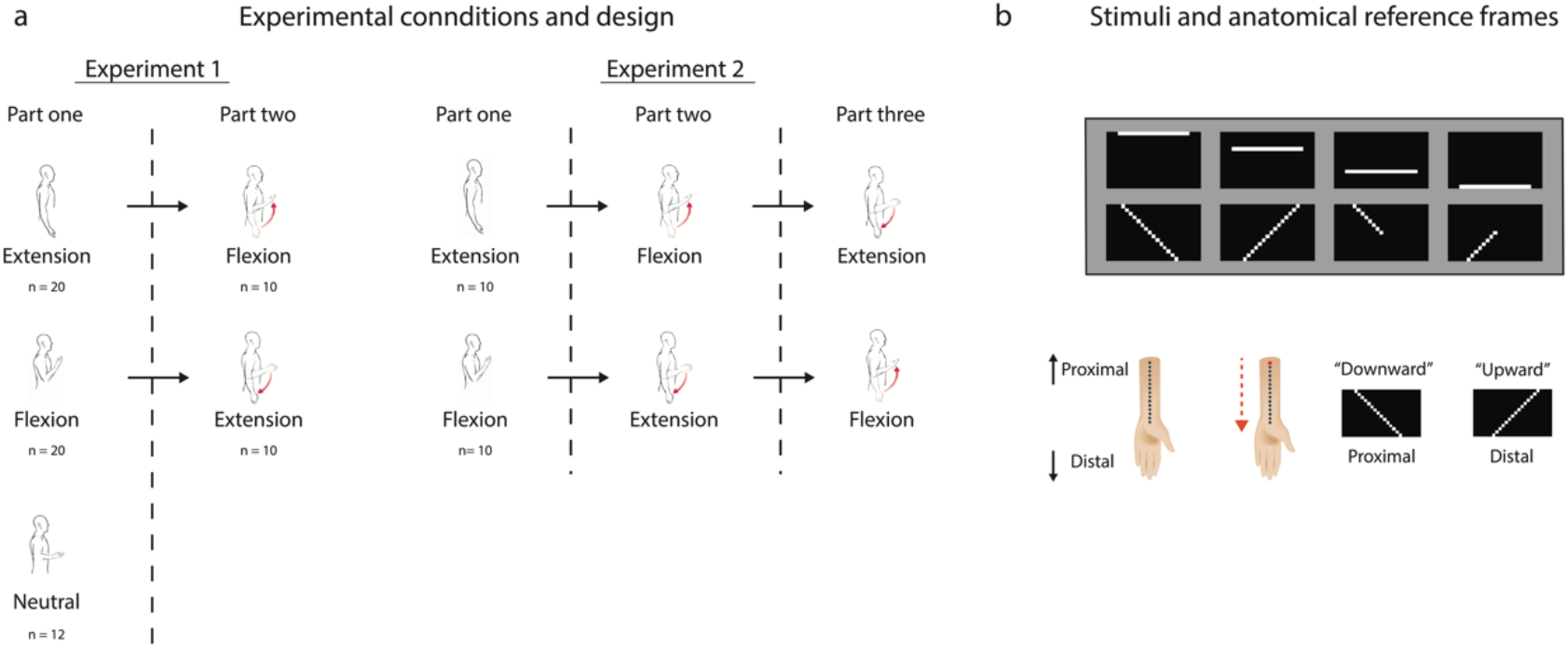
Experimental design. Participants in experiment 1 (**a**) were randomly allocated to one of three arm postures. At the end of part one, 20 participants were asked to switch their posture from Flexion to Extension or vice versa. Participants in experiment 2 were randomly assigned either to the Flexion or Extension posture and had to switch postures twice between blocks. 2D visual images (**b**) were transformed to vibrotactile stimulations on the inner arm. Responses on the stimulus’s orientation or location were interpreted based on the anatomical landmarks of the arm, resulting in a proximal or distal selection. The same stimulus can thus be interpreted as different reference frames according to the participant’s response.

### Experiment 1

#### <u>Part one</u> (n = 52)

In the Neutral posture (n = 12) participants showed no inclination (Fig. 2a.) on the group level for one coordinate over the other (W^+^ = 46, p = 0.1467), with 51.7 [23.5%, 79.9%] (mean, 95% confidence interval) of responses matching the distal selection (i.e., up location/orientation towards the wrist). In the Flexion posture (n = 20, Fig. 2a.), the average distal response was 95.3 [91.7%, 98.9%] that significantly differed from chance (W^+^ = 210, p = 0.00008). In the Extension posture (n = 20), 89.5% [79.7%, 99.3%] of responses matched a proximal selection (Fig. 2a.), with a significant difference compared to chance (W^+^ = 202.5, p = 0.00028).

**Fig 2.**
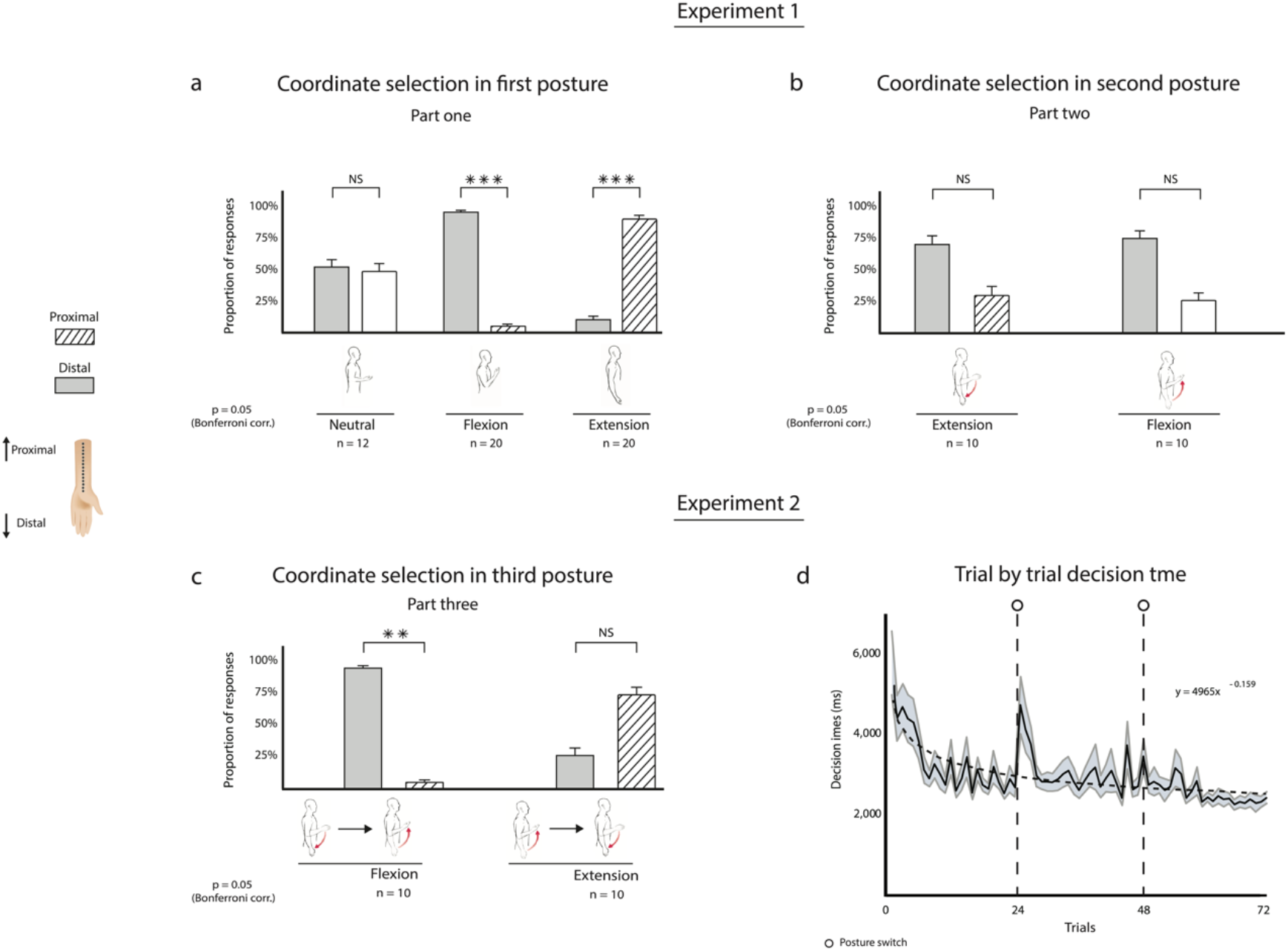
Group results (**a+b**) for distal and proximal responses in experiment 1. In part one (**a**) participants significantly chose a distal preference in the Flexion condition, a proximal preference in the Extension condition, and did not significantly choose neither in the Neutral condition. The difference in participants’ behavior among the different conditions highlights the implicit effect of posture on reference frame selection. After switching postures (**b**) participants in both the Flexion and Extension conditions did not show a significant preference for either a proximal or a distal selection. Earlier adaption of reference frames primed participants’ remapping after the posture switch. In part three of experiment 2 (**c**), following two posture switches, participants return to their initial posture assumed in part one with similar behavioral results. Group level decision times (**d**) tended to decrease across experimental parts. A fitted curve shows saturation in learning at about 60 trials, with little apparent effects of posture switching on the overall learning process.

#### <u>Part two</u> (n = 20)

Participants were first assigned either the Flexion or Extension posture in part one, and were then instructed to switch to the opposite posture. Following the posture switch, participants’ preferences in coordinate selection diverged from part one (Fig. 2b.). On the group level, for the Flexion posture (n = 10), distal responses after the switch were 74.4% [49.8%, 97.8%] with no significant difference when compared to chance (W^+^ = 42.5, p = 0.063). Likewise, in the Extension posture, participants did not show a preference after the switch with 30% [0.035%, 0.565%] of responses in line with a proximal coordinate selection (W^-^ = 12, p = 0.114).

### Experiment 2

In our second experiment, we replicated parts one and two of experiment 1 while recording participants’ (n = 20) decision times and adding another posture switch and trial block (part three).

#### Part one

For the Flexion posture (n = 10), the average distal responses were 93.6% [88.1%, 99.9%] that significantly differed from chance (W^+^ = 55, p = 0.0051). For the Extension posture (n = 10), 73.7% [49.1%, 98.3%] of responses matched a proximal selection with no significant difference compared to chance (W^-^ = 17, p = 0.284, for more information see subject-level analysis).

#### Part two

Average distal responses in the Flexion posture after the switch were 59.96% [31.2%, 88.8%] with no significant difference compared to chance (W^+^ = 34, p = 0.492). Similarly, there was no significant preference in coordinate selection after the switch for the Extension posture, with 34.3% [7.78%, 60.8%] of responses in line with a proximal coordinate selection (W^-^ = 22.5, p = 0.389).

#### Part three

After switching postures again, participants returned to their initial posture (either Flexion or Extension) with similar behavioral results (Fig. 2c.). The proportion of distal responses in the Flexion posture was 94.6% [91.2%, 98%] (W^+^ = 55, p = 0.0025). In the Extension posture, average proximal responses were 73.8% [0.5, 0.976] with no significant difference compared to chance (W^+^ = 39.5, p = 0.12).

#### Decision times

Decision times were collected for all 20 participants across the three experimental parts (Fig. 2d.). The average decision time in part one was 3413.42 [3250, 3580] milliseconds (ms), which dropped to 3098.06 [2950, 3250] ms in part two, with a further reduction to 2534.51 [2440, 2630] ms in the third block. To better understand the learning process and its related learning curve, we fitted a power-law function to the graph. The function has a slope of - 0.159 and intercept of 4965, Kolmogorov-Smirnov goodness of fit analysis resulted in no significant differences between the model and the data (R^2^ = 0.5481, p = 0.072). From the fit, we can predict some saturation in the learning process after approximately 60 trials. In contrast to the overall learning curve trend, there is a single significant increase in decision times after the first trial following the first posture switch (n = 20, p = 0.001).

### Subject-Level Analysis

To understand the reference frame choices during the tactile remapping process before and after switching, we analyzed participants’ responses from both experiments on each part. We asked whether a participant held one preference over the other compared to chance (Normal approximation to Binominal, X ~ B {n = 16-24, p = q = 0.5}, FDR corrected). All 72 participants showed a clear and significant preference (n = 72, a = 0.05) in part 1 (Fig. 3a.). For the Flexion posture all 30 participants took a distal reference frame, while for the Extension posture 27 out of 30 participants adopted the proximal one. In the Neutral posture, six participants showed a consistent proximal preference, and the other six a distal one. In part two, responses from 39 out of 40 participants passed the FDR correction (n = 40, a = 0.05) with a significant consistent preference (Fig. 3a.). 14 participants who switched from Extension-to-Flexion adopted a distal reference frame (switched coordinates) and 6 adopted a proximal one (maintained the same coordinates). For those who switched from Flexion-to-Extension, 13 adopted a distal reference frame (maintained the same coordinates) and 6 a proximal preference (switched coordinated). In sum, out of the 39 participants, 17 participants changed their reference frame after the switch, which indicates a remapping into external or trunk-centered reference frames, while 22 participants maintained their initial reference frame which indicates a remapping anchored to the anatomy of the inner arm. In part three, all 20 participants’ responses passed the FDR correction as they all reverted to the preferences selected in part one (Fig. 3a.). 10 participants adopted a distal reference frame in the Flexion condition. In the Extension condition 8 participants adopted the proximal reference frame and other two the distal one. These individual preferences show that the insignificant group result for the Extension condition in part three (Fig. 2d.) does not stem from participants’ inability to select and hold a reference frame.

**Fig 3.**
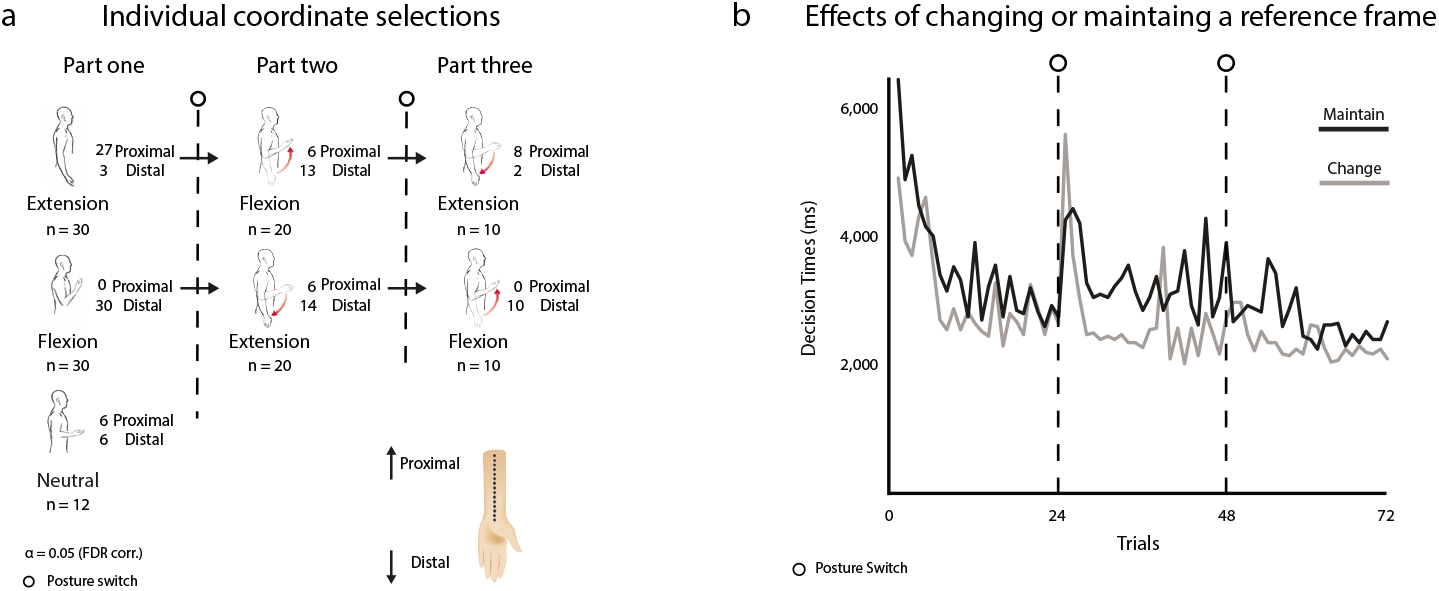
Individual coordinate preference (**a**) for all three parts aggregated on both experiments. To test for the strength and consistency of preferences, each participant’s responses were compared to chance and then corrected for multiple comparisons. All participants passed the correction for part one (n = 72), 39 passed for part two (n = 40), and all 20 passed for part three (n = 20). Post-hoc analysis of average decision times in experiment 2 (**b**) plotted separately for participants who maintained their preference after switching postures (n = 13) and participants who flip their perception of the y-axis (n = 7). The difference between the graphs appears constant throughout the experiment, even prior to the first posture switch.

To further investigate the process of changing or maintaining a reference frame, we conducted a post hoc analysis between decision times of participants who changed their reference frame after switching (External or trunk-centered, n = 7) and those who maintained their previous reference frame (arm-centered n = 13). We observed that participants who changed their reference frames have shorter overall decision times (Fig. 3b.). To examine the differences between the groups we conducted an ANCOVA on the regressions’ line coefficients. The difference in slopes were not significant between the two groups (df = 1, F = 2.01, p = 0.16) suggesting a similar learning curve for both groups. However, the intercept of participants who maintained their reference frame was significantly higher than those who changed reference frames (n = 72, p = 0.0001).

## DISCUSSION

The current study investigated the role of proprioception in tactile remapping by measuring the effects of arm posture on reference frame selections, and the extent to which they are affected by switching postures. We found that participants’ initial selection of reference frames is highly dependent on their posture, and was not anchored to a specific anatomical location on the inner arm, such as the wrist or the elbow. The remapping destination was to another body part, such as the trunk or face, which reflected an egocentric reference frame, or otherwise anchored to the external environment in an allocentric reference frame^1,4,9,10,17^. However, the neutral posture could neither prompt a trunk-centered nor an external reference frame as the arm is positioned perpendicularly to the trunk and to the up-down coordinates of the room. However, participants’ responses were individually consistent, and reference frames were easily adopted even in this ambiguous spatial position. This behavior might be related to the specific body part in use (i.e., a body-part to which we are trained to select a reference frame even in ambiguous spatial conditions), or due to the lack of guiding visual inputs that could cause confusion.

The ability to adopt reference frames independent of posture was further corroborated in part 2, where participants switched between opposing postures. According to the sensorimotor contingencies theory, switching postures should facilitate an adaptation period while new sensory information is integrated with stored body representations^13,16^. The cognitive cost in adapting to new postures is described as a multisensory integration problem that requires updating stored representations with new sensory information from many modalities. In line with the predictions of this theory, our results show that the selection of reference frames was strongly primed by the previous posture. About half of the group kept a reference frame that is centered on the trunk or external by reversing their perception of the vertical axis. In contrast to these predictions, the other half adopted a new reference frame that is based on the anatomy of the inner arm, which is invariant to changes in the arm’s posture. Most importantly, we observed little cost for either adopting a new or maintaining a reference frame. Participants exhibit a strong consistency in their responses after changing postures while their decision times are unaffected and continue to follow a general learning curve trend (save for the first trial after the switch). Although participants’ initial reference frame is trunk/face-centered or external, many of our participants switch to a reference frame that is centered on the anatomy of the inner arm. Moreover, our post-hoc analysis hints that participants who chose the inner arm as a reference may sport shorter decision times, even when considering trials that occurred before the switch. The latter result suggests that differences in decision times between the groups couldn’t be specifically linked with a posture switch, but perhaps associated with a more general behavior. The small and uneven number of subjects per group and the lack of differences in the learning rate between them calls for a more direct investigation by future studies.

Considering the weighing scheme model of sensorimotor contingencies^2^ in the context of our findings, the ability to select multiple reference frames with little cognitive costs follows an extreme instance where all options are weighed equally. It is possible that while the initial choice of reference frames is implicit, switching posture in the absence of vision forces participants to make an explicit choice. This is further corroborated by the results from the second posture switch, where all participants maintained their previous reference frame affirming an explicit choice. Taken together, our results show that top-down modulation can easily nullify low-order proprioceptive cues when choosing between reference frames, and that previously stored representations can be abstracted from current sensory inputs and spatial tasks. We suggest that the lack of visual inputs is the primary reason for the behavior. Vision is essential in forming body representations and has been widely reported as dominant over competing inputs from other modalities^2,19,25,26^. For example, crossing effects in temporal order judgments are substantially decreased when participants are blindfolded but the same manipulation has little effect on the weighing of different reference frames in congenitally blind adults^27–29^. Moreover, vision and touch share a combined multisensory object representation, which is formed by inputs from both modalities^30^. Visual cues thus act both as facilitators for body representations but also act as a disturbance to maintaining a stored body representation that was tactually derived. As our participants are blindfolded, vison could not override the changes in proprioceptive signals, revealing the contribution of proprioception to the coordinate transformation process. Proprioception is a unique sensory modality, and though much is known about its physiology, it remains a somewhat esoteric sensory modality. While vision is an exteroceptor identified with a known sensation, proprioception is an interoceptor that, for the most part, is not consciously perceived^24^. It is a perception of the self that results from motor actions taken and initiated by the self, and can thus be predicted. As such, the sensory consequences of arm movement could be anticipated, and they might interfere less with higher body representations. In conclusion, the present study demonstrates that top-down modulations can nullify new proprioceptive information during the process of tactile remapping, ultimately confirming that the weight of proprioceptive information during spatial tasks is considerably weaker compared to that of visual information.

## METHODS

### Participants

A total of 72 healthy participants took part in two experiments (age 30.8 ± 10.8, average and standard deviation; 44 females; 64 right-hand dominant). 52 participated in experiment 1 (age 32.4 ± 11.8; 31 females), and a further 20 took part in experiment 2 (26.5 ± 5.7; 13 females). Sample sizes were first chosen on the basis of first-level analysis, such that the number of overall trials is sufficient for a normal approximation which allows for a standardized t-test. However, in some cases the number of participants per condition allowed only a non-parametric second-level analysis, and further samples could not be collected due to the coronavirus crisis. No participants or samples were excluded from the study. The experiments were approved by the Interdisciplinary Center Herzliya (IDC) ethics committee, all participants signed an informed consent form and were paid for their participation in the study. The two experiments in this study were performed in accordance with relevant guidelines and regulations set by the IDC institutional ethics committee.

### The “Tactile Glove” – device description

The “Tactile Glove” is a custom-built Sensory Substitution Device (SSD) that conveys visual information from a 2D image into vibrotactile stimulations. The glove consists of 15 standard 8-millimeter diameter coin vibration motors which are placed in a row on the participant’s inner arm, another vibrator on the index finger acts as a precursor. Each actuator is supplied with a five Volt logic and interfaced with a Data Acquisition Module (iUSBDAQ-U120816, *HYTEK Automation*). An accompanying algorithm (written in C#) down-samples 2D images to a 15-by-25-pixel grayscale image, with white pixels denoting objects (e.g. lines or shapes). The binary image is temporally scanned from left to right, column by column using a sweep-line approach. For each white-colored pixel detected in a column, an actuator simultaneously vibrates on the inner arm. This procedure results in the image’s *Y*-axis represented by the spatial location on the arm, and the *X*-axis represented by timing (e.g., parts of the image that are heard sooner are positioned more to the left). Each stimulus comprises of a 300ms precursor cue, a short 100ms pause, and another 150ms spent on each column of the image. Participants had no prior familiarity with the device, the algorithm, nor with any other SSD.

### Stimuli and postures

The stimuli consisted of 8 black and white images: four images depicted horizontal lines and another 4 depicted diagonal lines (see Fig. 1b. for some examples of the stimuli). Three postures were used as experimental conditions (Fig. 1a.): a) Arm Extension, with the palm facing outward in full supination; b) Arm Flexion, with participants’ elbow leaning on an adjacent table, palm facing toward the face; c) An in-between position, named a Neutral posture, with the arm placed on the table and the palm facing the ceiling. We visually verified throughout the sessions that participants maintain their postures during experimental blocks.

### Procedure

We conducted two separate experiments of image localization, focusing on the perceived elevation (up-down) coordinate of the presented stimuli. Blindfolded participants were fitted with the device on their dominant hand, and received a short introduction about the device and experimental process, followed by two introductory pre-test stimuli. Importantly, no information was given on the way the algorithm conveys information on the y-axis. Participants had to report the orientation or spatial position of the stimuli. For horizontal line stimuli (Fig. 1b.), the question was “*Is the stimulus located on the upper or lower part of the image?*”, and for the diagonal line stimuli (Fig. 1b.), “*Does the stimulus have a downward or upward slope?*”. The experimenter did not provide any feedback to participants’ responses.

Experiment 1 included 16-24 randomized trials in each trial block (11:27, CI [10:55, 12:00] minutes, average experimental time and confidence interval). For each trial, the stimulus was repeated three times with 200 milliseconds interstimulus interval, followed by a verbal response from the participant. In part one, 52 participants were assigned to the Neutral (n = 12), Extension (n = 20), or Flexion posture (n = 20). In part two, 20 participants who performed the Flexion and Extension conditions were asked to switch their posture before completing another block of trials with the same task. To reduce implicit biases, participants were told that switching arm postures is necessary to eliminate fatigue (Fig. 1a.).

In experiment 2, participants were asked to respond with keyboard strokes as soon as possible, using their non-stimulated hand. To minimize the bias towards the keys’ physical location, participants were randomly assigned to different key stroke combinations. In part one, 20 participants were randomly assigned to the Extension (n = 10) or Flexion postures (n = 10). Participants were asked to switch their postures twice (Fig. 1a.), at the end of part one and of part two. Self-initiated responses resulted in shorter block durations (7:47, CI [7:13, 8:23] minutes) in comparison to the first experiment

## AUTHOR CONTRIBUTIONS

O.Y developed the Sensory Substitution Device and programed its accompanying code as well as the experiments’ graphic interfaces and data collection schemes. O.Y, G.B, and B.H contributed to the study design. O.Y and G.B performed the data collection, O.Y analyzed the data, interpreted the results, and produced all the figures. O.Y drafted the manuscript, G.B, B.H, D.F, and A.A provided important revisions and approved the draft for submission. A.A provided the funding for the study.

